# A Pilot Study on Assessing Auditory Masking Using Auditory Steady-State Responses

**DOI:** 10.1101/2025.11.06.686909

**Authors:** Anna Sergeeva, Preben Kidmose

**Affiliations:** Department of Electrical and Computer Engineering, Aarhus University, Denmark

**Keywords:** auditory masking, electroencephalography, auditory steady-state response

## Abstract

Auditory masking is important in the characterization of human hearing and hearing impairment. Traditionally, masking is assessed through behavioral methods, witch requires active participant engagement. This study investigates the potential of using Auditory Steady-State Response (ASSR) to assess auditory masking, enabling masking assessment without requiring active participation.

ASSRs were measured in response to a 40-Hz amplitude-modulated probe signal with and without the presence of a masker. The probe signals were 1/3-octave band-width Gaussian noise centered at 891 and 1414 Hz (center frequency, CF) and presented at 10, 20, 30, and 40 dB above individual behavioral masking thresholds (MT). The masker was lowpass Gaussian noise (cut-off 707 Hz) presented at 65 and 85 dB SPL (masker level, ML).

The ASSR amplitude increased with presentation level (PL) and decreased in the presence of a masker, confirming a masking effect on ASSR. At 65 dB ML, ASSRs did not differ between center frequencies when probe signals were presented relative to MT, suggesting a simple relationship between MT and ASSR. At 85 dB ML, an effect of CF was observed, suggesting that the relationship between MT and ASSR is more complex than initially anticipated, and involving all the experimental parameters (CF, PL, and ML).

## 1. INTRODUCTION

Auditory masking is a fundamental concept in audiology, describing the phenomenon where the perception of one sound (the probe-signal) is influenced by the presence of another sound (the masker). It is typically assessed through behavioral tests, where individuals are tasked with detecting the probe signal. The behavioral masking threshold refers to the minimum level of a probe signal allowing behavioral detection of the probe in the presence of a masker. Behavioral masking thresholds are commonly studied in psychoacoustic experiments and serve as the basis for models widely employed in hearing aid fitting [1]. Psychoacoustic models of masking have described frequency-specific masking effects and the critical band theory, which explains masking in terms of auditory filters and their bandwidths [2]. However, behavioral measurements are inherently subjective and depend on the participant’s attention, motivation, and cognitive state. Furthermore, significant variability in masking thresholds exists among individuals, particularly those with hearing loss [3]. This variability creates challenges for accurately predicting and compensating for masking effects in hearing aid algorithms.

Auditory Steady-State Response (ASSR) is widely used in clinical practice for hearing assessment, particularly in estimating frequency-specific hearing thresholds in individuals unable to perform behavioral tasks. ASSR reflects the neural system’s entrainment to the stimulus, and is typically measured in response to an amplitude modulated stimulus [4, 5]. This neural entrainment can be influenced by external factors, such as the presence of a masking sound. Masking affects the signal-to-noise ratio (SNR) of the auditory stimulus, and increased masking reduces SNR, resulting in decreased ASSR amplitude.

Evidence of the effects of masking on ASSR and other electrophysiological measurements has been reported in various contexts. Several studies have shown that masking noise applied contralaterally can suppress ASSR amplitudes [6–8]. Furthermore, contralateral noise, even at levels that do not cause significant behavioral threshold elevation, has been shown to significantly elevate the thresholds of the 40-Hz ASSR [9]. Similar effects of masking have been observed in other auditory measures, such as the Auditory Brainstem Response (ABR). For instance, the latency of wave V in the ABR increases with rising levels of broadband white noise masker, while the amplitudes of both wave I and wave V decrease [10].

Masking effects on the ASSR can be considered as an undesirable factor that negatively impacts its estimation. However, the masking effects can also be utilized to study the masking phenomenon itself. Unlike behavioral methods, this approach enables an objective assessment of masking effects without requiring active participation from the subject. To our knowledge, ASSR-based estimation of masking thresholds has not been previously explored.

This study examines the masking effect on the ASSR and its relationship to the behavioral masking threshold. The masker is a low-pass noise signal presented at two intensity levels. There were two probe signals, each a one-third octave amplitude-modulated noise, centered at different frequencies above the masker’s cutoff. Each probe was presented at four levels relative to the behavioral masking threshold. This experimental design allows us to quantify the masking effect on the ASSR as a function of presentation level and spectral distance from the masker.

In designing this study, we hypothesized that ASSR amplitude would increase monotonically with probe intensity, enabling masking threshold estimation using a methodology similar to ASSR-based hearing threshold estimation. Additionally, we expected the masking effect on the ASSR to be independent of the probe signal’s center frequency, provided the probe levels were set relative to the behavioral masking threshold.

By exploring the relationship between behavioral masking threshold and ASSR, this study aims to enhance our understanding of physiological masking mechanisms and assess the potential of ASSR as an objective tool for estimating masking thresholds.

## 2. METHOD

### 2.1 Subjects

Six participants (4 women, mean age: 33.2 years) with normal hearing thresholds (*<*20 dB HL for octave frequencies between 500 and 4000 Hz) and no history of hearing disorders were included in the study. Behavioral hearing thresholds for each ear were measured using the ascending method as outlined in ISO 8253-1:2010 [11]. This study was conducted as a preliminary pilot investigation to refine experimental procedures and assess feasibility. Participants were staff or students from our department who voluntarily agreed to participate based on informed consent.

### 2.2 Measurement setup

EEG was recorded concurrently from four scalp electrodes and 16 ear electrodes. The scalp electrodes were placed at left (M1) and right (M2) mastoids, Fpz and AFz according to the 10-20 EEG electrode system [12], and attached to the skin using custom designed electrode holders made of silicone with double adhesive pads. Prep pads were used to clean the skin prior to the attachment of the scalp electrodes. A small amount of gel (Electro-Gel, Electro-Cap International, Inc., USA) was applied on the scalp electrodes. The data recorded from ear-electrodes were not used in this study.

The EEG recordings were acquired with a sampling rate of 1000 Hz by a 32 channel portable MOBITA EEG amplifier (TMSi B.V., The Netherlands). The recording electrodes were Ag/AgCl electrodes with a diameter of 4 mm [13]. The ground electrode was placed on the neck using a disposable wet gel electrode (Ambu WS, Ambu A/S, Denmark). The auditory stimuli were presented to the test subjects using insert earphones (3M E-A-RTONE for ABR, 50 Ohm, 3M, USA) via an RME soundcard (Fireface UC, RME, Germany) with a sampling frequency of 48 kHz. Synchronization of the auditory stimuli and EEG recordings was achieved using a trigger signal generated by the soundcard. The trigger signal, periodic at 0.1 Hz, was sent to the EEG amplifier’s trigger input via a g.TRIGbox (g.tec medical engineering GmbH, Austria).

### 2.3 ASSR stimuli and recordings

The probe signal consisted of 40-Hz amplitude-modulated (AM) 1/3-octave Gaussian noise (GN) signals with center frequencies of 891 Hz and 1414 Hz. The masker was low-pass GN with a cut-off frequency of 707 Hz. Both the probes and the masker were filtered in the frequency domain. Initially, the GN signals in its entirely length were transformed to the frequency domain using a Discrete Fourier Transform (DFT). A window function was then applied to set frequencies outside the band of interest to zero. Finally, the complex spectrum was transformed back into the time domain using the Inverse DFT. This filtering method ensures a filter characteristic with steeper slopes than the auditory-filter bank. The masker was presented at levels of 65 and 85 dB SPL, while the stimuli levels were set at 10, 20, 30, and 40 dB above individual behavioral masking thresholds (MT). Stimuli were presented diotically. A schematic representation of the paradigm is shown in Figure 1.

**Figure 1.**
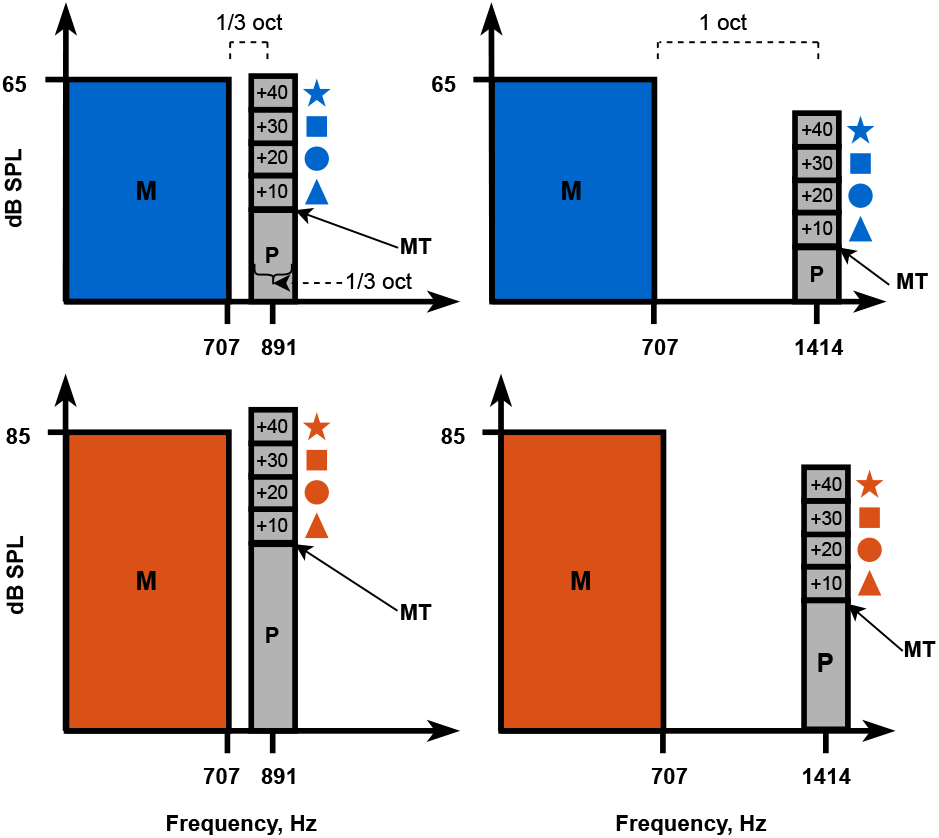
Schematic representation of the paradigm. The masker (M) was presented at levels 65 dB SPL (blue) and 85 dB SPL (orange). The probes (P) were presented at levels10 dB (triangles), 20 dB (circles), 30 dB (squares) and 40 dB (stars) above the behavioral masking threshold (MT), which were determined for each center frequency and masker level.

Behavioral masking thresholds were determined for each masker level and center frequency (500-ms 1/3-octave GN centered at 891 and 1414 Hz served as probe-signals) using a three-alternative forced-choice paradigm with a 1-up, 3-down adaptive procedure. The initial step size was 5 dB. After four reversals, the step size was reduced to 2 dB and the test proceeded until 12 reversals were reached. The threshold was calculated as the average of the last 8 reversals. A computer monitor displayed a response box with three buttons representing the stimulus intervals in each trial. During stimulus presentation, the buttons were highlighted successively in sync with the corresponding intervals. The subjects were instructed to press the button corresponding to the interval containing the probe and received immediate feedback indicating whether the response was correct or incorrect.

ASSR measurements were conducted for each combination of masker level, probe level and center frequency. In the presence of a masker, EEG was recorded for 20, 20, 10, and 10 minutes for stimuli presented at 10, 20, 30, and 40 dB above MT, respectively. Similarly, for reference without masker, EEG was recorded for 10, 10, 5, and 5 minutes at 10, 20, 30, and 40 dB above MT, respectively. Longer recording times were used at lower stimulus levels to account for the lower ASSR amplitudes, ensuring sufficient data to achieve a significant response.

The total recording time was 6 hours, distributed over three days. On the first day, recordings were conducted at a center frequency of 891 Hz with two masker levels and four probe levels, totaling eight recordings (2 hours). The same procedure was followed on the second day for a center frequency of 1414 Hz, also resulting in eight recordings (2 hours). On the third day, recordings were performed without the masker (16 trials, 2 hours). Within each recording day, stimuli were presented in randomized order.

Both behavioral tests and EEG recordings took place in a double-walled sound-attenuating room. During EEG recordings, participants were seated in a comfortable chair and instructed to relax but remain awake and to have open eyes. They were offered the option to watch a silent, subtitled movie of their choice to avoid becoming drowsy during the experiment.

### 2.4 Data analysis of EEG

Analysis of the EEG data was performed offline after the recordings were completed. ASSRs were estimated from the average of the mastoids with reference to Fpz. The data was band-pass-filtered using an 8th order, zero-phase filter with passband between 10 and 100 Hz and Butterworth characteristic. A notch filter was applied to remove 50 Hz line noise. The trigger signal was used to split the dataset into epochs of 10 seconds length and the epochs were averaged using weighted averaging [14].

The averaged epoch was transformed into the frequency domain by means of a DFT. The ASSR was determined as the amplitude at the modulation frequency bin, and the background noise was calculated as the root-mean-square in the frequency band ±6 Hz relative to the modulation frequency (excluding the modulation frequency) [4].

Statistical significance of the ASSR amplitude was determined based on an F-test [15]. The F-ratio was calculated as the ratio between the power at the modulation frequency and the average power in the frequency band ±6 Hz relative to the modulation frequency (excluding the modulation frequency). An F-ratio with a p-value ≤ 0.05 was regarded as statistically significant. Only statistically significant ASSRs were included in the subsequent statistical analysis.

### 2.5 Statistical analysis of the recorded ASSR

Differences between ASSRs in the presence and absence of the masker were evaluated using a paired permutation test [16] with 1000 permutations. For the statistical analysis of ASSRs in the presence of the masker, mixed-effects models were employed in R (lme4, version 1.1.35.5, Rstudio R-4.4.2) [17]. The model included *Center Frequency* (*CF*: 891 and 1414 Hz), *Masker Level* (*ML*: 65 and 85 dB SPL), *Presentation Level* (*PL*: 10, 20, 30 and 40 dB MT) and all their interactions as fixed effects. *Subject* (*S*) was included as a random effect to account for inter-subject variability. Statistical significance of the fixed effects was assessed through model reduction based on likelihood ratio test using the anova function in R. Post hoc analyses were conducted using estimated marginal means (EMMs) computed with the emmeans R package [18], with Bonferroni correction applied to adjust p-values for multiple comparisons.

## 3. RESULTS

Table 1 presents the mean and standard deviation of behavioral masking thresholds (MT) in dB SPL (relative to 20 *µ*Pa) across six subjects for each masker level (ML) and carrier frequency (CF). For comparison, it also includes the mean and standard deviation of behavioral thresholds measured without a masker using the same procedure.

**Table 1.** Mean (standard deviation) of the behavioral masking thresholds in dB SPL across the subjects for each masker level and without the masker.

| Center Frequency | Masker Level |  |  |
| --- | --- | --- | --- |
|  | 65 dB SPL | 85 dB SPL | Without |
| 891 Hz | 29.8 (4.7) | 55.7 (2.3) | 7.6 (4.1) |
| 1414 Hz | 18.8 (2.9) | 39.5 (3.5) | 9.6 (2.2) |

For each subject, ASSR was recorded in 32 different conditions (2 CF × 4 PL × 2 ML × with/without masker). Six subjects were included in the study, yielding a total of 192 ASSR recordings, of which 2 recordings were not significant. Thus 190/192, corresponding to 99 % of all recordings, were significant.

Figure 2 illustrates the relationship between ASSRs to different stimuli in the experimental design, both with and without the masker. The data is shown for each individual subject followed by the grand average (only significant data points are shown). The figure reveals a reduction in ASSR amplitude in the presence of the masker, particularly at lower presentation levels. This difference diminishes as the presentation level increases.

**Figure 2.**
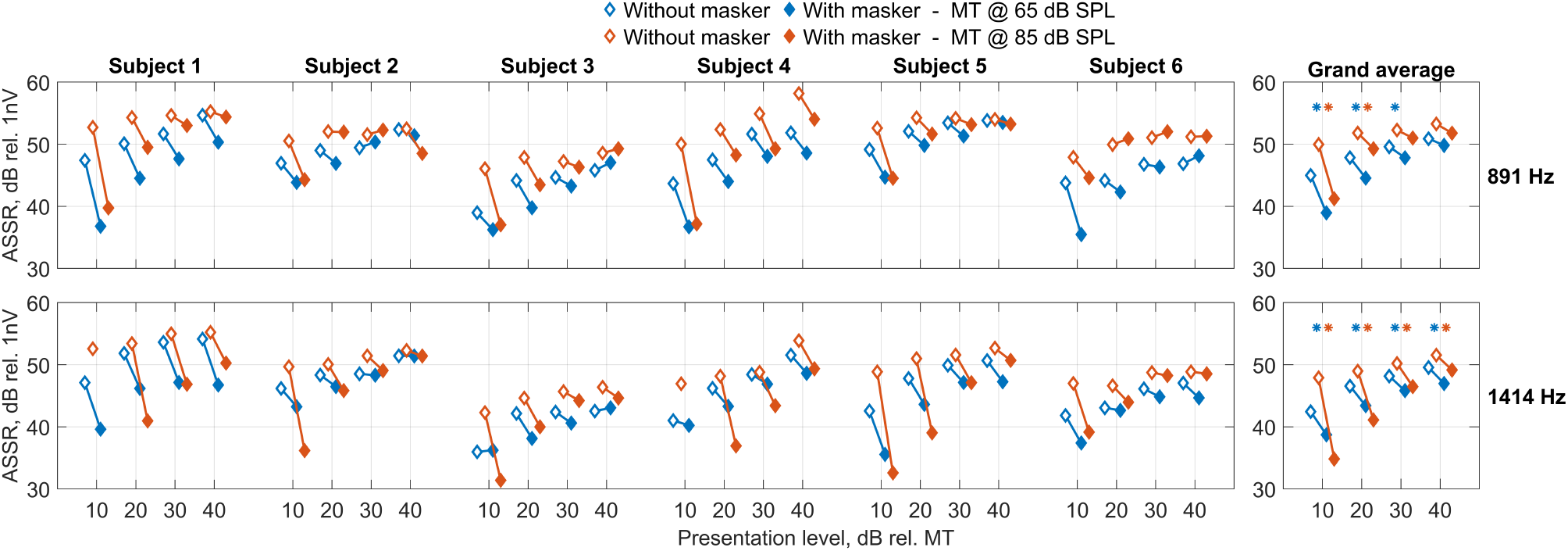
ASSR as a function of the presentation level for each CF (891 Hz – upper subplots, 1414 Hz – lower subplots) and masker level (blue for 65 dB SPL, orange for 85 dB SPL). Data is shown with (filled rhombs) and without (empty rhombs) the masker. Left: individual subject data; right: grand average across subjects. Asterisks indicate significant difference between ASSR with and without the masker (blue for 65 dB SPL, orange for 85 dB SPL).

Table 2 provides p-values from permutation tests comparing ASSRs with and without the masker. Significant differences are marked with asterisks in the grand average plots shown in Figure 2.

**Table 2.** p-values from paired permutation tests, comparing ASSR with and without the masker for each masker level, center frequency and presentation level.

| Center Frequency | Masker Level | Presentation Level |  |  |  |
| --- | --- | --- | --- | --- | --- |
|  |  | 10 dB MT | 20 dB MT | 30 dB MT | 40 dB MT |
| 891 Hz | 65 dB SPL | 0.001 | 0.001 | 0.049 | 0.303 |
| 891 Hz | 85 dB SPL | 0.001 | 0.049 | 0.227 | 0.110 |
| 1414 Hz | 65 dB SPL | 0.013 | 0.001 | 0.001 | 0.045 |
| 1414 Hz | 85 dB SPL | 0.001 | 0.001 | 0.001 | 0.001 |

Figure 3 illustrates the relationship between ASSR and center frequency (891 and 1414 Hz) for stimuli presented with masker at two levels (65 and 85 dB SPL) across various presentation levels relative to the masking threshold. The data points represent the same ASSR measurements with masker shown in Figure 2, but they are visualized in a different way. Data is shown both for individual subjects and as the grand average across all subjects. The figure demonstrates that when the probe signal is presented relative to the masking threshold, ASSR amplitudes remain relatively stable across center frequencies when the masker level is 65 dB SPL. In contrast, for masker level of 85 dB SPL, a noticeable reduction in ASSR amplitude is observed at 1414 Hz compared to 891 Hz.

**Figure 3.**
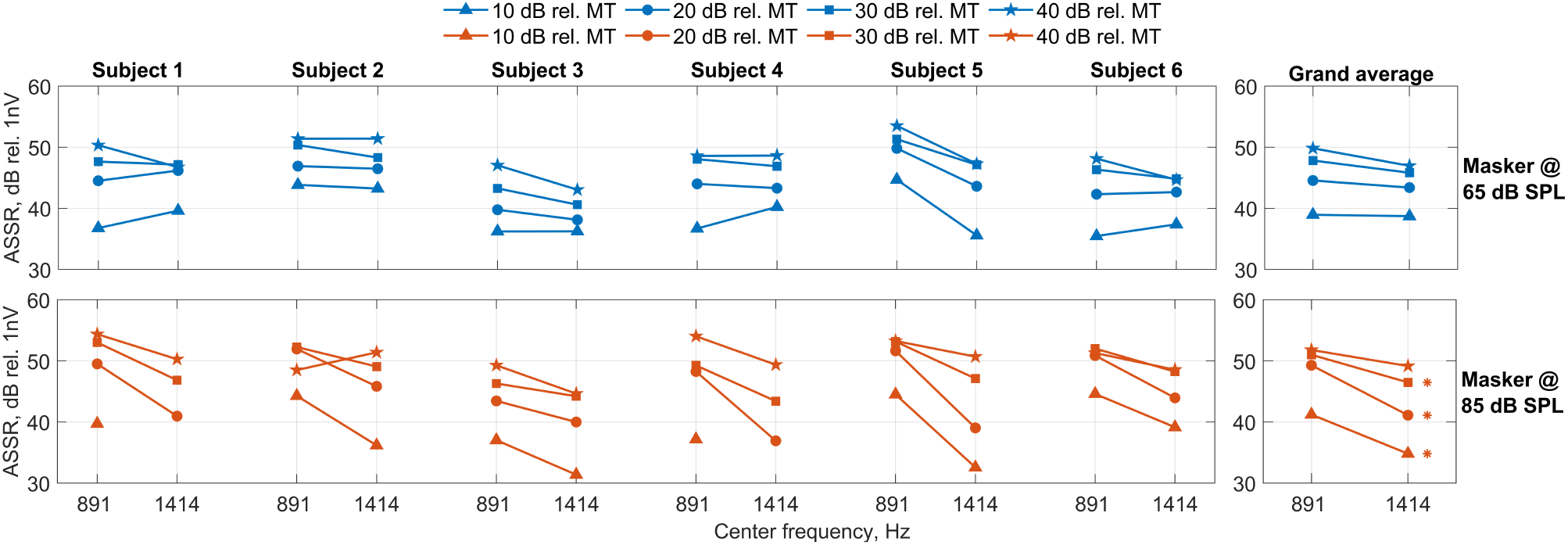
ASSR as a function of the CF for stimuli presented 10 dB (triangles), 20 dB (circles), 30 dB (squares) and 40 dB (stars) relative to masking threshold. Data is shown for masker levels of 65 dB SPL (blue, upper subplots) and 85 dB SPL (orange, lower subplots). Left: individual subject data; right: grand average across subjects. Asterisks indicate significant difference in ASSR amplitude between 891 Hz and 1414 Hz.

A full mixed-effects model was fitted to the ASSRs with the masker: *ASSR* ∼ *CF + ML + PL + CF:ML + CF:PL + ML:PL + CF:ML:PL + (1*|*S)*

Residuals met model assumptions for normality (Shapiro-Wilk test: w = 0.989, p = 0.678) and homogeneity of variance (Bartlett’s test: *χ*^2^(15) = 22.996, p = 0.084). Likelihood ratio tests during model reduction revealed a significant three-way interaction (*CF:ML:PL, χ*^2^(3) = 12.1, p = 0.007), indicating that the model could not be simplified. Table 3 presents the post hoc analysis of estimated marginal means comparing ASSRs at center frequencies 891 Hz and 1414 Hz. Significant reductions at 1414 Hz are marked by red asterisks in Figure 3.

**Table 3.** Post hoc analysis of estimated marginal means comparing ASSR at different center frequencies for each masker level and presentation level, SE - standard error of mean.

| Masker Level | Presentation Level | ASSR<sub>891Hz</sub> – ASSR<sub>1414Hz</sub> | SE | t.ratio | p-value |
| --- | --- | --- | --- | --- | --- |
| 65 dB SPL | 10 dB MT | 0.234 | 1.25 | 0.188 | 1.000 |
| 65 dB SPL | 20 dB MT | 1.159 | 1.25 | 0.930 | 1.000 |
| 65 dB SPL | 30 dB MT | 2.008 | 1.25 | 1.611 | 0.881 |
| 65 dB SPL | 40 dB MT | 2.871 | 1.25 | 2.304 | 0.186 |
| 85 dB SPL | 10 dB MT | 6.458 | 1.40 | 4.604 | <.001 |
| 85 dB SPL | 20 dB MT | 8.167 | 1.25 | 6.553 | <.001 |
| 85 dB SPL | 30 dB MT | 4.522 | 1.25 | 3.628 | 0.003 |
| 85 dB SPL | 40 dB MT | 2.635 | 1.25 | 2.114 | 0.147 |

## 4. DISCUSSION

The behavioral masking thresholds presented in Table 1 align with previous empirical findings and established auditory system theories [1]. Initially, we observe a a clear increase in the threshold when a 65 dB masker is present compared to the without-masker condition. This confirms that the masker has an effect and that the measured threshold is a masking threshold rather than a hearing threshold. At both center frequencies (891 Hz and 1414 Hz), thresholds were higher at a masker level of 85 dB SPL than at 65 dB SPL, confirming that masking thresholds increase with masker level. Notably, consistent with established masking theories, a 20 dB increase in masker level resulted in a more than 20 dB increase in masking threshold. Additionally, in this study, where the masker was low-pass filtered noise with a 707 Hz cutoff frequency, the masking threshold was higher at 891 Hz than at 1414 Hz. This finding supports the auditory filter bank theory, which suggests that masker penetration decreases as the detection band is further away from the masker bands [1, 2].

Figure 2 illustrates the ASSR amplitudes as a function of presentation level for each center frequency (CF) and masker level (ML), with data shown both with and without the masker. The results indicate that ASSR amplitudes are generally lower in the presence of a masker compared to when a masker is absent, which is consistent with findings previously observed in the literature [6, 7]. This masking effect is particularly prominent at lower presentation levels, where the difference between ASSR with and without the masker is most pronounced. As the presentation level increases, the masking effect diminishes, and ASSR amplitudes with the masker approach those observed without it.

These findings are statistically supported by the paired permutation test results presented in Table 2. At 1414 Hz, significant differences were observed at all presentation levels for both masker levels. At 891 Hz, the effect was significant at 10, 20, and 30 dB MT for the 65 dB SPL masker, and at 10 and 20 dB MT for the 85 dB SPL masker, with no significant difference at higher presentation levels. This pattern further confirms that the masking effect is stronger at low probe levels and becomes less pronounced as the probe level increases. We hypothesize that, when the probe is near the masking threshold, the probe signal - and consequently, its modulation - is masked by the masker signal to a high degree. However, as the probe level increases, an increasingly larger part of its modulation surpasses the masking threshold, resulting in a stronger neural representation of the modulation.

Additionally, Figure 2 demonstrates a clear monotonic increase in ASSR amplitude with increasing presentation level, both in the presence and absence of a masker. This trend is consistent across subjects and frequencies. The observed increase in ASSR amplitude aligns with previous studies showing that higher presentation levels recruit a greater neural population, leading to stronger responses [19–21]. Although the presence of a masker reduces the ASSR amplitude, the relationship between ASSR and presentation level continues to exhibit a monotonic increasing pattern as a function of presentation level. This suggests that ASSR can serve as an objective measure for predicting behavioral thresholds. Specifically, in the absence of a masker, the threshold corresponds to the hearing threshold, whereas in the presence of a masker, it corresponds to the masking threshold.

Figure 3 illustrates ASSR amplitudes as a function of center frequency (891 Hz vs. 1414 Hz) for different presentation levels relative to masking threshold (MT) and masker levels (65 dB SPL and 85 dB SPL). The figure shows that with a 65 dB SPL masker, ASSR amplitudes remain relatively stable across frequencies at all presentation levels. In contrast, with an 85 dB SPL masker, ASSR amplitudes at 1414 Hz are notably lower than at 891 Hz, with this effect being most pronounced at lower presentation levels. This pattern suggests a complex interplay between masker level, center frequency, and presentation level, indicating that physiological masking is not simply a linear function of these parameters.

A mixed-model reduction procedure confirmed this complexity by revealing a significant interaction between masker level, presentation level, and center frequency. To further examine this interaction, post hoc analyses were conducted. At 65 dB SPL, no significant differences were found between 891 Hz and 1414 Hz across all presentation levels, suggesting that there is the same relationship between ASSR and behavioral masking threshold across probe frequencies.

However, at 85 dB SPL ASSR amplitudes at 1414 Hz were significantly lower than at 891 Hz for presentation levels of 10, 20, and 30 dB MT, indicating a stronger physiological masking effect at the higher masker level. Interestingly, no significant difference was observed at 40 dB MT. One possible explanation is that at this high presentation level, ASSR amplitudes at 891 Hz may have reached a saturation point, reducing the observable difference between the two frequencies. Previous studies have reported that ASSR amplitudes tend to saturate at levels above 80 dB SPL [22–24].

## 5. CONCLUSION

This study was based on the hypothesis that the ASSR amplitude depends on the probe’s presentation level relative to the behavioral masking threshold. While the empirical findings strongly support such relationship, they also reveal a more complex interaction involving the masker level and the probe signal’s center frequency relative to the masker’s cut-of frequency. Future studies will further explore these interactions.

## 6. ACKNOWLEDGMENTS

This study is supported by Center for Ear-EEG at Department of Electrical and Computer Engineering, Aarhus University, Denmark.

